# CDK12 and CDK13 suppress distinct intronic polyadenylation sites

**DOI:** 10.64898/2026.05.15.725461

**Authors:** Molly Hulver, Andrew Polevoda, Kai-Lieh Huang, Eric J. Wagner, Paul L. Boutz

## Abstract

Cyclin-dependent kinases CDK12 and CDK13 are RNA polymerase II C-terminal domain Ser2 kinases that regulate transcriptional elongation and co-transcriptional RNA processing. Loss-of-function (LOF) mutations in these kinases are observed in multiple cancers and are characterized by increased intronic polyadenylation (IPA), which generates prematurely terminated transcripts and can give rise to novel intronic peptides. To investigate the distinct contributions of CDK12 and CDK13 to suppressing early transcriptional termination, we modeled selective kinase inhibition in OVCAR4 ovarian cancer cells using THZ531, a dual CDK12/13 inhibitor. A CRISPR-engineered CDK12 C1039S mutation conferred THZ531 resistance, enabling functional distinction between CDK12 and CDK13 LOF conditions.

Using Poly(A) Click-seq (PACseq) across a six-point inhibitor dose curve, we identified thousands of IPA sites with distinct dose-response behaviors, allowing us to classify CDK12- and CDK13-dependent subsets. Oxford Nanopore long-read sequencing confirmed premature termination events and revealed full-length isoform switches, demonstrating how IPA alters coding potential by truncating protein domains and producing novel, intronically encoded peptide sequences. In parallel, integration of long-read transcriptomes with mass spectrometry data identified peptides derived from IPA isoforms, confirming that intronic sequences can be translated.

Together, these findings establish distinct roles for CDK12 and CDK13 in maintaining transcript integrity, demonstrate that IPA transcripts are not only abundant but also translationally competent, and provide evidence for a mechanistic link between transcriptional dysregulation and the generation of novel peptide isoforms in CDK12/13-deficient cancers. These results highlight a potential therapeutic vulnerability through the exploitation of IPA-derived peptides as tumor-specific neoantigens.

## Introduction

Transcriptional programs must be carefully orchestrated to enable cells to respond to changes in the environment. Modulation of the transcriptional machinery on a short time scale (minutes to hours) is carried out through the dynamic phosphorylation/ dephosphorylation of the RNA Polymerase II C-terminal Domain (RNAPII CTD) heptapeptide repeats^1–5^. Specific phosphorylation events alter the association of the polymerase with other components found within the pre-initiation complex and at the promoter-proximal pause site, and coordinate the complexes that promote traversal of the chromatin environment and the processing of the nascent RNA through splicing and cleavage and polyadenylation (CPA)^6,7^. The most abundant and broadly functional of these modifications is phosphorylation of serine 2 (pSer2) of the RNAPII CTD, which is carried out by three cyclin-dependent kinases in vertebrates: CDK9, which releases the polymerase from the promoter-proximal pause, and CDK12/CDK13, paralogous kinases that both maintain pSer2 through the elongation phase of transcription as well as phosphorylate other target proteins involved in transcription, RNA processing, and RNA decay^8–12^.

During transcription elongation, CDK12/13 both promote polymerase processivity. Depletion of CDK12 kinase activity, through loss-of-function (LOF) mutations, genetic depletion, or small-molecule inhibitors, results in increased termination at early polyadenylation sites (PAS) often present within the introns of genes (intronic polyadenylation, IPA)^12,13^. Because many genes encoding the homologous recombination (HR) machinery contain multiple, highly CDK12-dependent IPA sites, loss of CDK12 activity profoundly downregulates HR repair of double-stranded DNA breaks^12^. Tumors with biallelic CDK12 LOF mutations thus phenocopy other mutations in HR genes (such as BRCA1/2) conferring a ‘BRCAness’ phenotype. The resulting genomic instability is thought to drive further tumor progression; hence CDK12 is a ‘master regulator’ of HR and functions as a tumor suppressor^14–17^.

CDK13 shares ∼95% amino-acid identity with CDK12 in its kinase domain, but the remainder of the protein is highly divergent from its homolog^18^. Promotion of transcriptional elongation through pSer2 is a shared function between CDK12 and CDK13^19–21^. Germline heterozygous mutations in the kinase domain of CDK13 result in congenital disease affecting cardiac and cognitive development^22^. Originally identified in melanomas, similar mutations are found broadly in multiple tumor types and are invariably heterozygous LOF. Depletion of CDK13 activity also results in an increased accumulation of prematurely-terminated RNA (ptRNA) transcripts, similar to that observed in CDK12 LOF. However, the mechanism is distinct as CDK13 phosphorylates Ser275 of the RNA binding protein ZC3H14—a component of the PAXT connection—which in turn recruits the nuclear exosome to decay the ptRNA target^22,23^. Thus the accumulation of transcripts terminating at IPAs occurs through an increase in stability, rather than production, of the ptRNAs. It has been observed that the IPA-terminated transcripts that increase in response to either CDK12 or CKD13 depletion are largely distinct groups. However, the small molecule drug THZ531, which is a potent and specific inhibitor of both kinases, is equally effective on both and thus cannot be used to distinguish which sets of transcripts are directly affected by either protein^24^.

Alternative polyadenylation has emerged in recent studies as a major determinant of transcriptome diversity^25^. A large fraction of human genes contain multiple PAS whose activity is regulated in a cell-type or environmentally dependent manner^25,26^. Often the canonical last exon of a gene will contain multiple PAS, in which case the coding region of the transcript is unaffected by which site is chosen, but the resulting 3’ UTR is different. This configuration is referred to as ‘tandem’ UTR PAS^27,28^. Proliferating cells have been observed to often favor the more proximal sites, shortening the 3’ UTR and eliminating binding sites for microRNAs and destabilizing proteins that are found in the distal PAS isoforms favored in quiescent cells^27,29–31^. Another common feature of genes is the presence of IPA sites found throughout the introns. When termination occurs at such sites, the coding sequence of the transcript is often truncated. Importantly, the use of an upstream IPA is a ‘zero-sum’ situation for the full-length transcript, so in suppressing the use of IPA sites one function of CDK12 is to promote the production of the complete mRNA transcript^13^. CPA is carried out by a large multi-protein complex that includes multiple RNA binding proteins that can recognize divergent elements surrounding the PAS and thus enabling a specific site to be up- or down-regulated in a controlled manner. How these sites and their surrounding sequence elements coordinate the CPA machinery and how CDK12 or CDK13 might modulate these activities to alter the production of different RNA isoforms is not well understood^4,6,19^.

Typically, 3’ end sequencing is used to provide deep coverage of the 3’ ends of polyadenylated transcripts, which are poorly represented in standard RNA sequencing libraries. Recent advancements in library preparation have used ‘Click’ chemistry to chemically attach adaptors to the RNA. PACseq is one such technique, which provides exceptional read depth at low per-sample costs^32^. However, despite the extraordinary depth that can be attained using these methods, the bioinformatic analysis of alternative polyadenylation is still highly non-standardized. A typical pipeline that is used for this type of analysis, PolyAMiner, is designed to detect the ‘tandem UTR’ type isoform switches, but is less usable for analyzing intronic polyadenylation^32^. Development of computational pipelines focused on this phenomenon would move the field forward substantially. Even more biologically relevant, long-read sequencing platforms such as Oxford Nanopore offer the capability of combining transcript structure with coding sequence production, which would enable the determination of the effects of early transcription termination on the encoded protein sequence, including those peptides encoded within introns which are normally removed and would not be expected to undergo translation. As with the alternative polyadenylation, there is little in the way of standardized methods to combine these powerful types of data.

In regards to the biological impact of CDK12 and CDK13 loss-of-functiohn (LOF), CDK12 biallelic loss-of-function (LOF) has been reported in ovarian, breast, and prostate cancers while CDK13 dominant negative mutations are found in melanomas^13,19–21,33–44^. Phenotypically, cases of CDK12 and CDK13 LOF result in increased IPA events where the polyA tail is added within an intronic region upstream of the canonical polyA site^12,13^. While the mechanism for how CDK12 LOF induces IPA is unknown, speculation can be made that its loss alters the kinetics of RNAPII and co-transcriptional processes. On the other hand, CDK13 dominant negative mutations likely disrupt the PAXT complex^22,45^. CDK13 phosphorylates the PAXT complex, or the polyA-tail exosome targeting complex, which degrades early terminated and polyadenylated mRNAs, including IPA isoforms. Thus, if CDK13 no longer activates PAXT, there is an accumulation of IPA transcripts^22,45^.

Both mammalian cell models and zebrafish models overexpressing mutant CDK13 are shown to express novel IPA mRNA transcripts not found in WT CDK13 models. Additionally, some of these transcripts are translated into novel proteins that contain an intronic peptide sequence which would typically be removed during splicing^22^. If a similar phenomenon occurs in patient tumors, this creates an exploitable therapeutic vulnerability: novel intronic peptide sequences could be targeted as neoantigens.

Antigens are peptides that have been processed within the cell and are then presented by MHCI proteins on the cell surface to be read by CD8+ T-cells^46,47^. T-cells will read the bound antigens and classify them as either self or non-self. Non-self antigens are then recognized as neoantigens and further trigger the immune response. Neoantigens can arise from invasive proteins found in the cell but can also be sourced from novel peptide sequences translated from mutated or not typically expressed endogenous mRNAs. Tumor immunotherapy treatments have been popularized to target neoantigens that come from mutated mRNAs that are unique to tumor cells and not found in a patient’s healthy cells^46,48,49^. The majority neoantigens being targeted by immunotherapy treatments arise from single nucleotide variant mutations and are not relatively abundant in the cell. In contrast, potential neoantigens that arise from tumor cells with either CDK12 or CDK13 LOF mutations would be expressed at much higher rates and be sourced from intronic regions of the mRNA rather than point mutations.

In this study, we have integrated multiple platforms to investigate the transcriptional changes in response to the CDK12/13 inhibitor, THZ531. Using genetic models to isolate the functions of CDK12 and CDK13, combined with a 6-point dose curve enabling high-resolution determination of the sensitivity of individual cleavage sites, we are able to distinguish multiple classes of IPA sites that are either CDK12 or CDK13 specific. These IPA sites are found in different functional groups of genes. Using long-read sequencing, we observe the effects of CDK12/13 dependent IPA usage on full-length protein-coding isoforms, demonstrating how the altered use of PAS sites in a gene can profoundly affect important protein-coding domains. Finally, using this novel transcriptomic data, we interrogate proteomics data from the inhibitor treatment to identify neo-peptides that are upregulated in response to CDK12/13 LOF and pointing toward possible targets for immunotherapy of tumors carrying CDK12/13 mutations.

## Results

### Differential sensitivity of ovarian cancer cell lines to THZ531 treatment

In order to model CDK12 and CDK13 LOF we attempted to generate point mutations in either CDK12 or CDK13 using CRISPR-Cas9 to create resistance to the inhibitor THZ531. THZ531 is a covalent inhibitor that binds to a cysteine residue outside of the kinase domain on both CDK12 and CDK13, so that when either protein is in its endogenously folded state, bound THZ531 physically blocks the kinase domain and prevents its phosphorylation activity^24^. Three ovarian cancer cell lines with no known CDK12 or CDK13 mutations were selected as potential candidates for THZ531 inhibitor resistance transfections. To determine which of these cell lines are sensitive to drug treatment, we treated them with 500 nM and 1µM of THZ531 along with vehicle alone (DMSO). We performed western blots from protein that was extracted 8 hours post-treatment and looked at the relative pSer2 levels of the RNAPII CTD. Each cell line had a distinct response to THZ531 (Fig.1A). OVCAR8 had minimal response even after 1µM of THZ531 as indicated by the relatively intense pSer2 band—indicating that OVCAR8 cells may exhibit resistance to the drug via metabolic or cellular efflux pathways. In contrast, CAOV3 cells were extremely sensitive to THZ531 treatment, demonstrating complete loss of pSer2 upon treatment with 500 nM THZ531. While decreased pSer2 is expected, total depletion suggests a possible off-target effect on CDK9, as it also phosphorylates the Ser2 residues of the heptapeptide repeat^50^. The final cell line we tested, OVCAR4, had a dose-response that best fit our experimental objectives, as both the 500 nM and 1 µM THZ531 treatments incompletely decreased pSer2 levels suggesting a specific inhibition of CDK12/13. Thus we selected OVCAR4 as the parental cell line to generate THZ531 resistance mutations.

**Figure 1:**
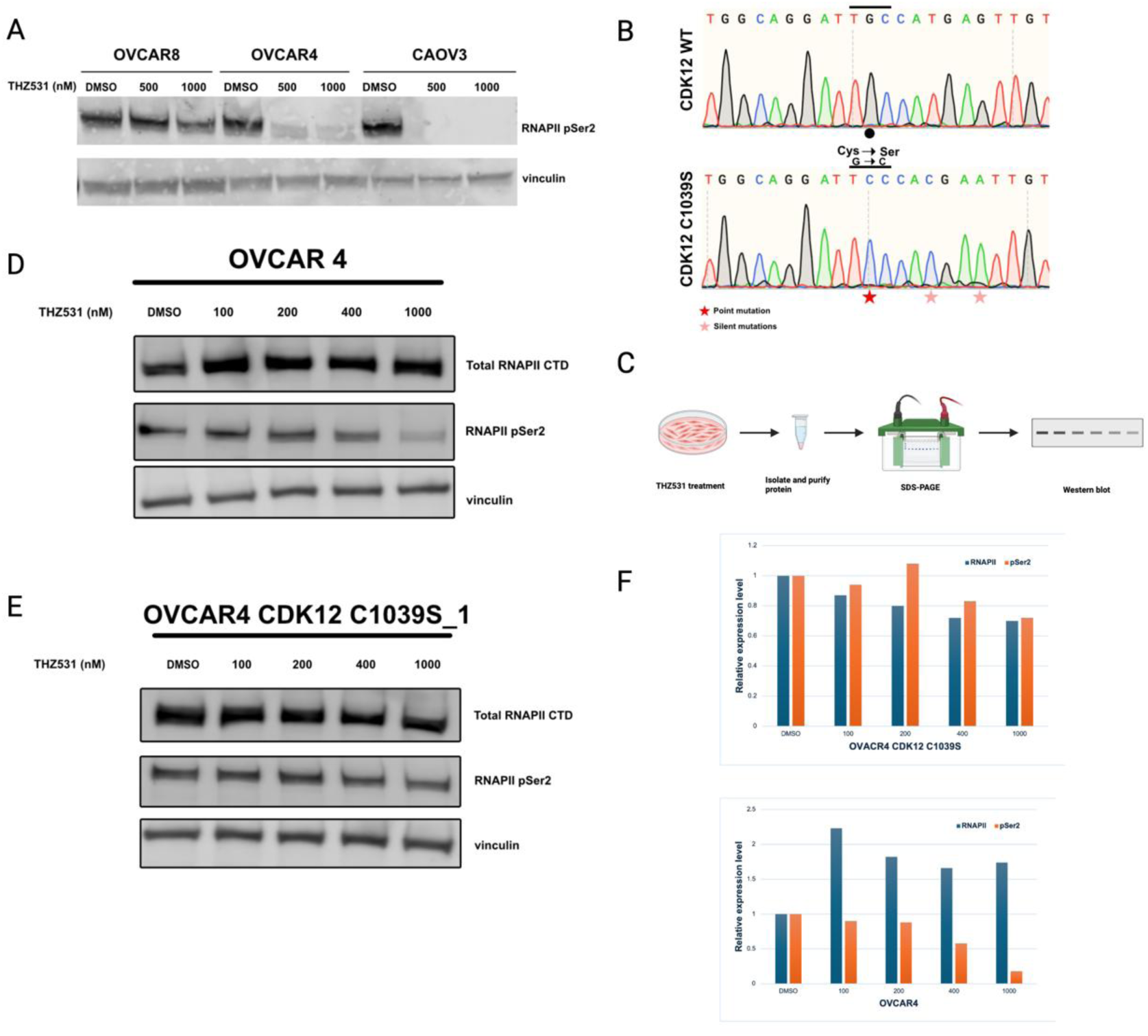
CDK12^C1039S^ mutation in OVCAR4 cells partially rescues depletion of CTD pSer2 after THZ531 treatment. A) OVCAR8, OVCAR4, and CAOV3 cells were treated with THZ531 and protein was blotted for pSer2 levels. B) Representative sequencing data CDK12 showing WT and the C1039S mutation, which is induced by a point mutation from guanosine to cytosine. Two silent mutations are also induced into the endogenous gene. C) A schematic of THZ531 treatment performed on OVCAR4 WT and C1039S cells. Cells were treated for 8 hours with THZ531, protein was isolated and purified, run on an SDS-PAGE gel, and then western blots were performed. D and E) Western blot of THZ531-treated OVCAR4 WT and C1039S cells. Total RNAPII levels remained steady upon increasing dosages. In WT cells, pSer2 levels dramatically increased upon increasing concentrations (D) but did not decrease to the same extent in CDK12 C1039S cells (E). F) Quantitative analysis of D and E with relative expression levels compared to DMSO-treated cells.

### CDK12 kinase domain mutants are resistant to THZ531

THZ531 forms a covalent bond with a cysteine residue in the CDK12 and CDK13 kinase domains, resulting in irreversible inhibition of their kinase activities^24^. The targeted residues are C1039 in CDK12 and C1017 in CDK13^24^. To generate inhibitor resistance mutants, we used an RNP-based CRISPR approach to direct an HDR template carrying a serine point mutation at each relevant cysteine, along with two silent mutations, to the cut site of both CDK12 and CDK13. We attempted to create two distinct cell lines: a C1039S CDK12 cell line and a C1017S CDK13 cell line. Following electroporation of the recombinant Cas9/sgRNA complexes with the HDR templates, clonal cell populations were isolated and genomic DNA sequenced. We identified three C1039S CDK12 OVCAR4 homozygous mutants, but no C1017S CDK13 mutants. Figure 1B shows Sanger Sequencing from one CDK12 homozygous mutant compared to WT CDK12: cysteine was mutated to a serine amino acid residue by changing the guanosine nucleic acid to cytosine. Two silent mutations were also introduced (T to G and G to A) to prevent the re-cutting of the DNA by Cas9. Although transfections of C1017S CDK13 were repeated multiple times, we did not identify any homozygous mutants, suggesting that mutations affecting the CDK13 kinase domain may be deleterious in OVCAR4 cells. Although the C1017S mutation in CDK13 is not expected to affect protein function in the absence of THZ531, it is possible that the mutation impacts basal kinase activity to a degree that cells carrying the mutation are at a disadvantage in terms of survival or proliferation.

### CDK12^C1039S^ mutation induces resistance to the dual CDK12/CDK13 kinase inhibitor

CDK12^C1039S^ OVCAR4 cells were treated with THZ531 for 8 hours at 100, 200, 400, and 1000 nM^12^. Protein was extracted and purified then run on an SDS-PAGE gel and western blots were performed. This workflow is illustrated in Figure 2.1C. Western blots were performed with antibodies recognizing either total RNAPII or pSer2-CTD (Figs.1D and E). While total RNAPII levels remained static over increasing THZ531 concentrations, pSer2 decreased in abundance in samples treated with higher THZ531doses in WT OVCAR4 (Fig.1D). In CDK12^C1039S^ mutant cells, pSer2 levels decreased but less so compared to WT CDK12 cells, indicating that mutant CDK12 is resistant to THZ531 and able to phosphorylate Ser2. The pSer2 and total RNAPII bands were quantified and relative abundance of pSer2 was compared to total RNAPII expression (Fig. 1F). For WT OVCAR4 cells pSer2 levels steadily decreased over increasing concentrations of THZ531. However, the mutant CDK12 cells exhibited no quantifiable decrease in pSer2 levels even at 1µM inhibitor treatment.

### 3’ end sequencing using PACseq and bioinformatic analysis of cleavage sites genome-wide

In order to identify 3’ cleavage sites genome-wide using a broad 6-point THZ531 dose response curve, we used ‘Poly-A Click seq’ (PACseq) to generate 36 independent replicate libraries in both wild-type and CDK12 ^C1039S^ mutant cells (Fig. 2A) which were sequenced on an Illumina NovaSeq. The PACseq libraries generate reads originating at the poly(A) tail or at genomically encoded poly(A) stretches within transcripts and spanning ∼150-200 nucleotides upstream of the cleavage site (Fig. 2B). We obtained approximately 1.1 billion reads, with 31.4 million reads on average per replicate. In order to take advantage of the high read depth and to make an unbiased analysis of the 3’ cleavage sites with respect to their location within genes, we developed a novel analysis pipeline (Fig. 2C). After aligning reads in a strand-specific manner with the genome, we required mapped reads to have a 3’ region that was ‘soft-clipped’ (i.e. did not align to the genome) and consisted of minimum of 6 nucleotides, all of which were adenosines such that the reads that passed this filter should have a minimum length polyA-tail that was not genomically encoded. We also eliminated reads terminating withing coding exons as we could not verify that these were *bona fide* cleavage sites. Lastly, we required that the sites have one of 8 consensus PAS sites (AATAAA, ATTAAA, etc) within 40 nucleotides upstream of the cleavage site. After filtering, we obtained ∼2.5 million unique 3’ cleavage sites within all replicates combined.

**Figure 2:**
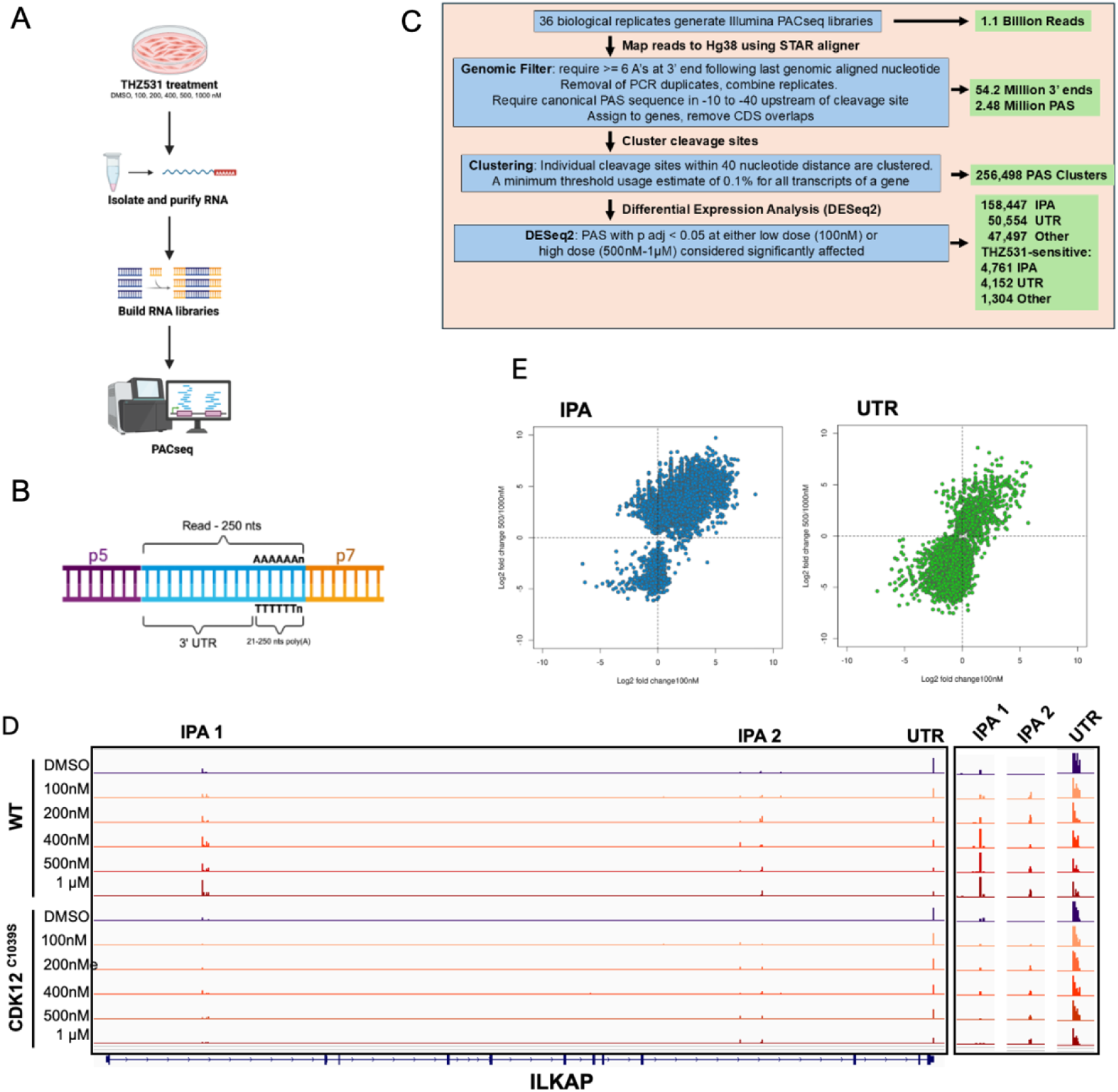
PACseq analysis of THZ531-treated WT OVCAR4 and CDK12 C1039S cells. A) Schematic of THZ531 treatment experiments. B) Schematic of prepared cDNA for PACseq analysis with 5’ and 3’ adapters. C) Schematic of bioinformatic pipline. Major processing steps (blue boxes), and the numbers of events identified (green boxes). D) Polyadenylation sites in ILKAP transcripts for all THZ531-treated samples. WT-treated cells exhibited increasing IPA site usage and decreasing UTR PAS usage. CDK12 C1039S mutant cells are almost entirely suppressed for IPA site usage. E) Scatterplot showing the log2-fold change in the 100nM THZ531 treatment vs. DMSO (x-axis) or 500nM+1000nM doses vs. DMSO (y-axis). IPA sites (left panel, blue) or UTR-associated sites (right panel, green) and filtered at a p adj < 0.05 are plotted. IPA sites generally increased with dosage, while more UTR sites decreased with dosage.

Because CPA-generated cleavage events often exhibit heterogeneity in the exact nucleotide position at which the cleavage occurs, we next clustered the sites that occurred within a short window (40 nucleotides). Clustering was performed at single-nucleotide resolution using the 3’-most mapped nucleotide of each read to determine the cleavage position. After step-wise clustering of each condition followed by re-clustering including the entire data set, we obtained a final genome-wide map consisting of 256,504 unique 3’ cleavage site clusters (Fig. 2C). Cleavage sites mapped to annotated sites within 3’ UTRs, within introns (IPAs), and other regions without known transcript structure (other, Fig. 2C). When we aligned the cleavage sites across drug doses within individual genes, we could observe many sites in which a dose-dependent increase in IPA sites occurred, at the expense of the distal UTR sites (Fig. 2D). Additionally, in many instances the increase in IPA site usage was blunted in the presence of the CDK12 C1039S mutation, while other sites were affected similarly in both wild type and mutant cell lines. For a number of sites, we observed that the increase in IPA site usage peaked at the 500nM concentration, suggesting the drug had achieved saturation at that concentration. For this reason, we combined replicates at the 500nM and 1uM doses in the differential expression analyses described below.

Using the genome-wide map of all detected PAS clusters, we next mapped and counted, at single-nucleotide resolution, each individual replicate at every drug dose and performed differential analysis using DESeq2. We performed separate low-dose (100nM) and a high-dose (combination of 500nM and 1uM) analyses in order to identify PAS exhibiting up- or down-regulation in response to the inhibitor, in both the wild-type and mutant cell line samples. We further annotated each individual cluster depending on its genomic location. Specifically, clusters falling within annotated introns of a gene were designated IPAs, those falling within annotated 3’ UTRs as UTR (further divided into the first occurring site (proximal) or those more downstream (distal), or single if only one site was present), and finally ‘other’ to indicate sites which did not meet either criterion. These would include sites falling outside of any annotated intron or UTR.

These analyses identified 4,761 IPA clusters that were significantly changed in expression in either low or high drug doses, or both (Fig. 2E). The majority of the IPA sites increased in usage upon drug treatment compared to control. We also identified 4,152 significantly changed UTR sites, and the large majority of these were downregulated in the drug treated cells, as well as 1,304 significantly changed ‘other’ sites. A comparison of the results using our novel pipeline and DESeq2 for differential expression with those obtained by another software package for alternative polyadenylation analysis, PolyAMiner, are ongoing (data not shown). The observations that IPA sites generally increase and UTR sites decrease are consistent with multiple previous studies. We next asked whether our 6-point dose response curve and exceptionally deep 3’ end sequencing data could enable us to identify direct CDK12- and CDK13-dependent IPA sites based on the precise inhibitor dose response of the individual clusters.

To perform genome-wide dose response measurements, we developed a novel hierarchical clustering method that allowed us to group clusters based on the relationship between different drug doses and resulting change in the fraction of transcripts within a gene that terminated at each PAS cluster. This analysis grouped the 4,761 IPA sites into 9 distinct clusters which could be grouped into 4 dominant metaclusters (Fig. 3A). Plotting the fraction of usage of each IPA site within its host gene, we could fit dose-response curves to each cluster and metacluster (Fig. 3B and Fig. 4A). These curves clearly indicated that metacluster 1 represents the set of IPA sites that exhibit a sigmoidal dose-response, with drug saturation occurring around 500nM; metacluster 2 had a parabolic curve, where peak increase in IPA site usage occurred around 200 nM and then began to decline at higher doses. Metacluster 3 exhibits a decline in usage from the control samples at each drug dose, while metacluster 4 is mostly unchanged across the dose regimen. The last cluster may represent sites whose host gene levels are changing significantly from the control to the low or high dose, but the fraction of usage of the site is relatively unchanged. This last group underscores the utility of our approach measuring the difference in the fraction of transcripts from a gene that terminate at each individual site, rather than treating each site as an independent entity within the genome-scale analysis.

**Figure 3:**
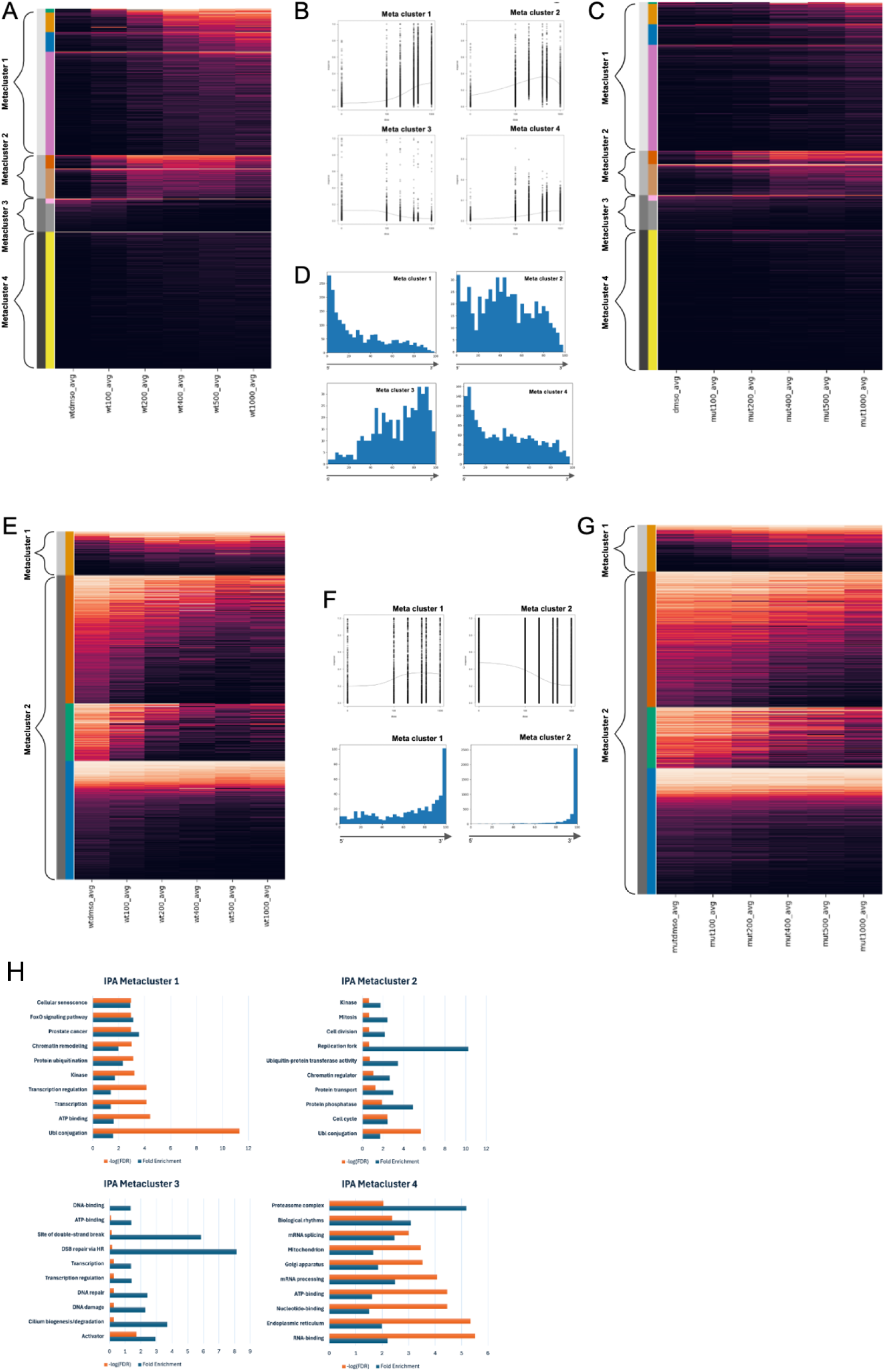
THZ531-treated cells undergo large-scale PAS usage changes within intronic and UTR regions. A) Heat map of IPA sites in THZ531-treated WT OVCAR4 cells. Affected genes are grouped into 9 clusters as indicated by the color-coded rows on the left, which are grouped into 4 metaclusters. The fraction of usage for each IPA site within its host gene is indicated by the color scale, at different drug doses (columns). B) THZ531 dose response curves of IPA sites grouped by meta cluster. C) Heat map of IPA sites in THZ531-treated OVCAR4 CDK12 C1039S cells grouped and arranged in the same order as in Figure 2.3A. D) Metagene frequency plots of IPA sites within normalized gene length bins, grouped by metacluster. E) Heat map of 3’ UTR sites in THZ531-treated WT OVCAR4 cells. Affected PAS are grouped into 4 clusters as indicated by the color-coded rows on the left, which are in turn grouped into 2 metaclusters. F) Dose response curves of the two UTR metaclusters (top) and UTR metagene frequency plots (bottom) grouped by the two metaclusters. G) Heat map of UTR sites in THZ531-treated OVCAR4 CDK12 C1039S cells grouped and arranged in the same order as in Figure 2.3E. H) Barplots indicating the fold enrichment and –(log10) of the p adj for the top ten gene ontology categories, grouped by each of the IPA metacluster containing genes.

**Figure 4:**
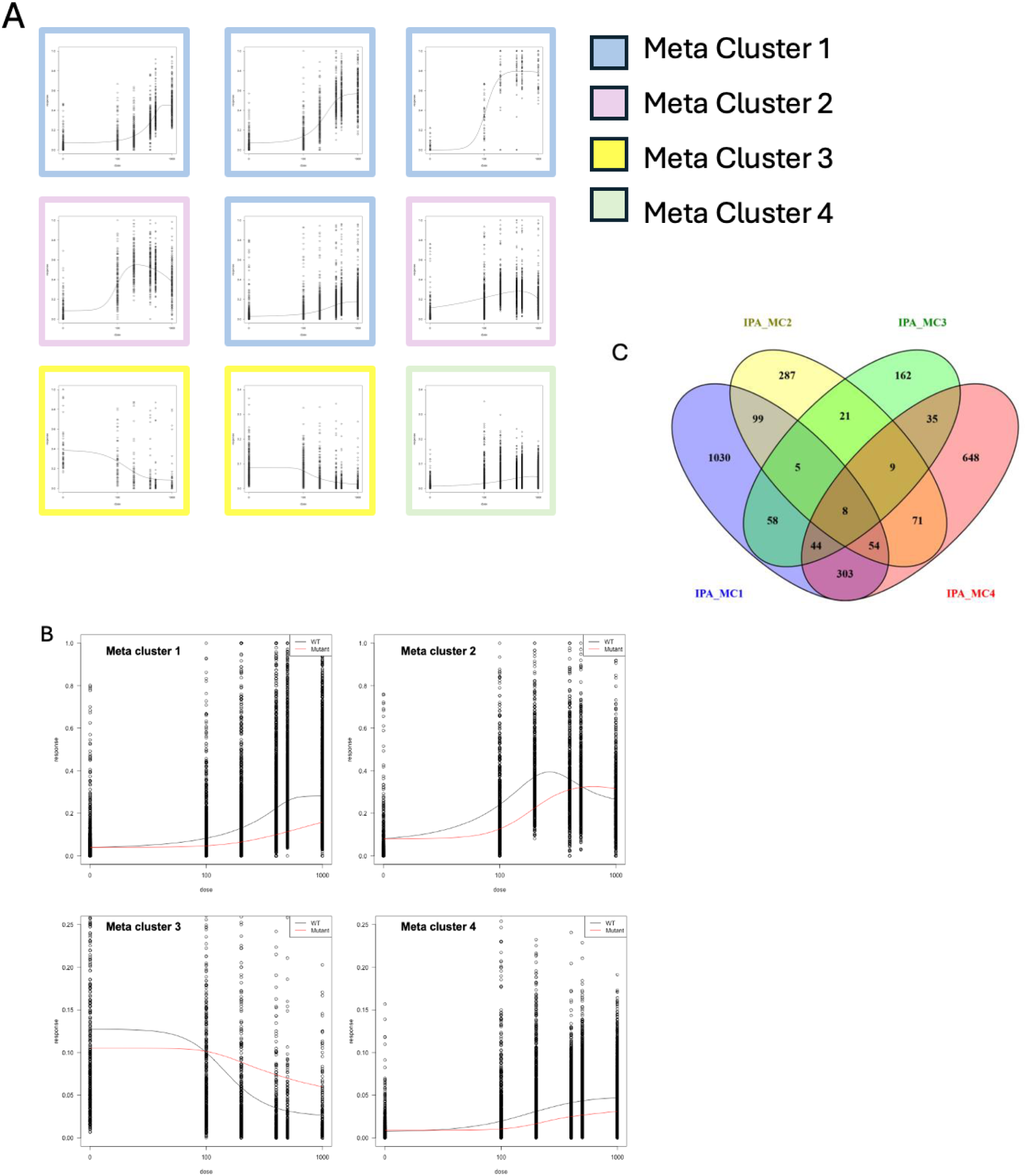
IPA metaclusters. A) The nine total IPA clusters dose-response curves from WT-treated cells and how they are sorted into the four metaclusters. The four clusters of metacluster 1 all follow a sigmoidal curve, metacluster 2 dose responses fit a parabolic curve, metacluster 3 responses decrease at higher THZ531 concentrations, and the single response curve of metacluster 4 remains largely unchanged. B) Overlaps of the dose-response curves of each metacluster of WT cells (black) overlapped with those in CDK12 C1039S cells (red). MC1 decreases in IPA usage over increasing concentrations of THZ531 in mutant cells, the MC2 response maintains the pattern but shifts slightly to the right. MC3 is similar in wild type and mutant, though with a less steep curve in the mutant, and MC4 remain mostly unchanged though may resemble metacluster 1. C) Venn diagram of IPA sites in the four IPA metaclusters. Each overlap indicates genes that have at least one IPA site in the indicated metacluster.

We next compared the effects of THZ531 treatment on the cells carrying homozygous CDK12 THZ531-resistance mutations (C1039S). As in the wild type cells, the usage fraction of each IPA site was plotted across the dose curves maintaining the same order of IPA sites as in the wild type heatmap (Fig. 3C). IPA sites within metacluster 1 were almost completely re-suppressed in the mutant cells, indicating that this metacluster consists of IPA sites whose usage is primarily affected by CDK12 kinase activity and not CDK13. At 1 µM, there was a small increase which may come from partial inhibition of mutant CDK12 by the drug at high concentrations. Metacluster 2 exhibited a slightly reduced sensitivity to drug dose, resulting in a shift of the dose response curve to the right. However, the parabolic response curve and the peak fraction of usage of these sites was mostly maintained, indicating that this cluster consists of sites that are mostly affected by the inhibition of CDK13 and not CDK12 (Fig. 3B). Metaclusters 3 and 4 were mostly unchanged in the mutant cells, though there are some sites that undergo increased usage in the mutant compared to wild type cells at the higher doses. These effects are seen clearly when the dose-response curves for wild type and mutant are overlayed (Fig. 4B).

Because termination at a given IPA site precludes the use of any other PAS, we considered the possibility that the dose response curves of metaclusters 2-4, rather than representing the intrinsic sensitivity of the corresponding PAS to inhibitor, could derive from some sites falling downstream of highly sensitive metacluster 1 sites. As the upstream site increases, the sites located downstream would decrease. We first performed a metagene analysis in which we binned the relative position of each IPA site within its host gene, and plotted a histogram for each metacluster (Fig. 3D). Metacluster 1 was highly skewed toward the 5’ end of genes, metacluster 2 spread more evenly throughout the gene body, and metacluster 3 frequency increased through the gene body, peaking at the 3’ end of genes. The distributions of the metaclusters are consistent with the hypothesis that the high sensitivity sites in metacluster 1, occurring early in the gene body, could be soaking up a large fraction of the transcripts from genes containing them, thus depleting downstream site usage as the drug dose increased. Metacluster 4 was most similar to metacluster 1, with strong enrichment at the 5’ ends of genes. However, when we determined the frequency with which IPA sites falling in each metacluster co-occurred within the same gene, we observed that the largest sets were the genes that had only one type of IPA site (Fig. 4C). Only about half of the sites within metaclusters 2-4 occurred in genes that also contained another type of IPA. The largest overlap was between metaclusters 1 and 4, which also shared a distribution pattern with enrichment toward the 5’ ends of genes. However, careful observation determined that often metacluster 4 sites could occur upstream of metacluster 1 sites. Altogether, these data indicate that positional effects cannot account for the different dose response curves of the metaclusters.

We next examined the effect of THZ531 treatment on PAS occurring within annotated UTRs. Overall, the majority of affected UTR sites were strongly downregulated (Fig. 3E). The sites formed four clusters and two metaclusters, with the much smaller metacluster 1 consisting of sites that increased in usage over the drug dose panel, and the large metacluster 2 comprising UTR associated PAS that decreased upon drug treatment. These trends could be observed when the fraction of usage values were plotted and a curve fitted to them (Fig. 3F). When we performed metagene analysis, metacluster 1 sites were spread throughout the gene body despite a strong 3’ localization, whereas metacluser 2 was mostly localized to the extreme 3’ end. These patterns likely suggest the presence of UTRs belonging to stable, upstream alternative last exons in metacluster 1 that could be upregulated by THZ531 treatment. In contrast, the decreasing metacluster 2 likely represents the bulk of genes in which an effect on one or more upstream IPA sites impacts the amount of full-length transcripts that are produced in the presence of the drug. In strong contrast to some of the IPA metaclusters, the predominant pattern observed with UTR sites was that the CDK12 resistant mutant cells were mostly unchanged compared to the wild type, though slightly less sensitive to downregulation at lower doses (Fig. 3G). Overall these trends suggest that CDK12 and CDK13 predominantly affect IPA sites. PAS located within distal UTRs are affected by the cumulative effects of upstream site usage triggered by inhibition of either kinase, but the effects on individual distal PAS are less likely to be a direct consequence of CDK12/13 inhibition.

Gene ontology analysis of the 4 IPA metaclusters grouped the functionality of the genes that were affected by IPA and further reinforced the hypothesis that these metaclusters represent differentially regulated gene sets (Fig. 3H). The top 10 significant clusters were graphed by FDR value and fold enrichment. The top hits within IPA metacluster 1were related to protein ubiquitination, transcription and its regulation, chromatin remodeling, cellular senescence, and prostate cancer. Given metacluster 1 is likely associated with CDK12 LOF which is found in prostate cancer, prostate cancer as one of the top gene ontology hits further confirms this conclusion. Metacluster 2 contains genes most associated with ubiquitination as well, but also includes the cell cycle (mitosis), and the replication fork. Interestingly, Metacluster 3 contains significant gene clusters associated with DNA repair and the homologous recombination repair pathway, which are associated with BRCAness phenotypes and have been previously associated with CDK12 LOF. Metacluster 4, which largely remains low and unchanged throughout increasing doses of THZ531 has significant clusters involved in mRNA processing, RNA binding, and specifically, the spliceosome. Further analysis of the gene clusters will explore these processes more deeply.

### Long-read sequencing defines coding sequences in alternatively polyadenylated messages

To elucidate the functional consequences of changes in alternative polyadenylation site usage upon THZ531 treatment, we sequenced wild type and CDK12 C1039S mutant cells treated with DMSO or low- or high-dose THZ531 using the Oxford Nanopore cDNA-PCR sequencing method. After mapping the reads and processing the data we obtained full-length transcript annotations and quantifications. With these data, we could observe the full-length transcripts produced under both control conditions and with THZ531 treatment, resulting in increases in transcripts ending at early termination sites (Fig. 5A). These full-length transcripts terminated at precisely the same nucleotide position indicated by the PACseq sites, and the polyA tails were visible within the reads (Fig. 5B). We used the analysis tool IsoformSwitchAnalyzeR to identify genes that exhibited significant changes in isoform usage between samples. An example of an isoform switch in the gene encoding Mediator subunit 27 is shown (Fig. 5C and D). The predominant full-length protein coding isoform (isoform 1) decreases from approximately 60% of the transcripts in the control samples to just over 20% in the THZ531 treated samples. The other full-length isoform that comprises most of the remaining MED27 transcript in the control samples also drops below 20%. In the THZ531 treated cells, isoforms terminating at multiple IPA sites early in the gene all increase significantly upon drug treatment (Fig. 5C,D). Genome wide, this tool identified ∼250 isoform switches involving alternative termination sites that reached significance (Fig. 5E). Notably, the read depth at which we obtained these results is significantly lower than the depth at which we could obtain PACseq reads. Additionally, multiple IPA sites within a single gene can each produce a unique isoform, the more of which are present in a gene, the more difficult it is for the isoform to reach a statistically significant value. With further refinement and increased read depth, the number of significantly detected isoform switches will likely increase substantially.

**Figure 5:**
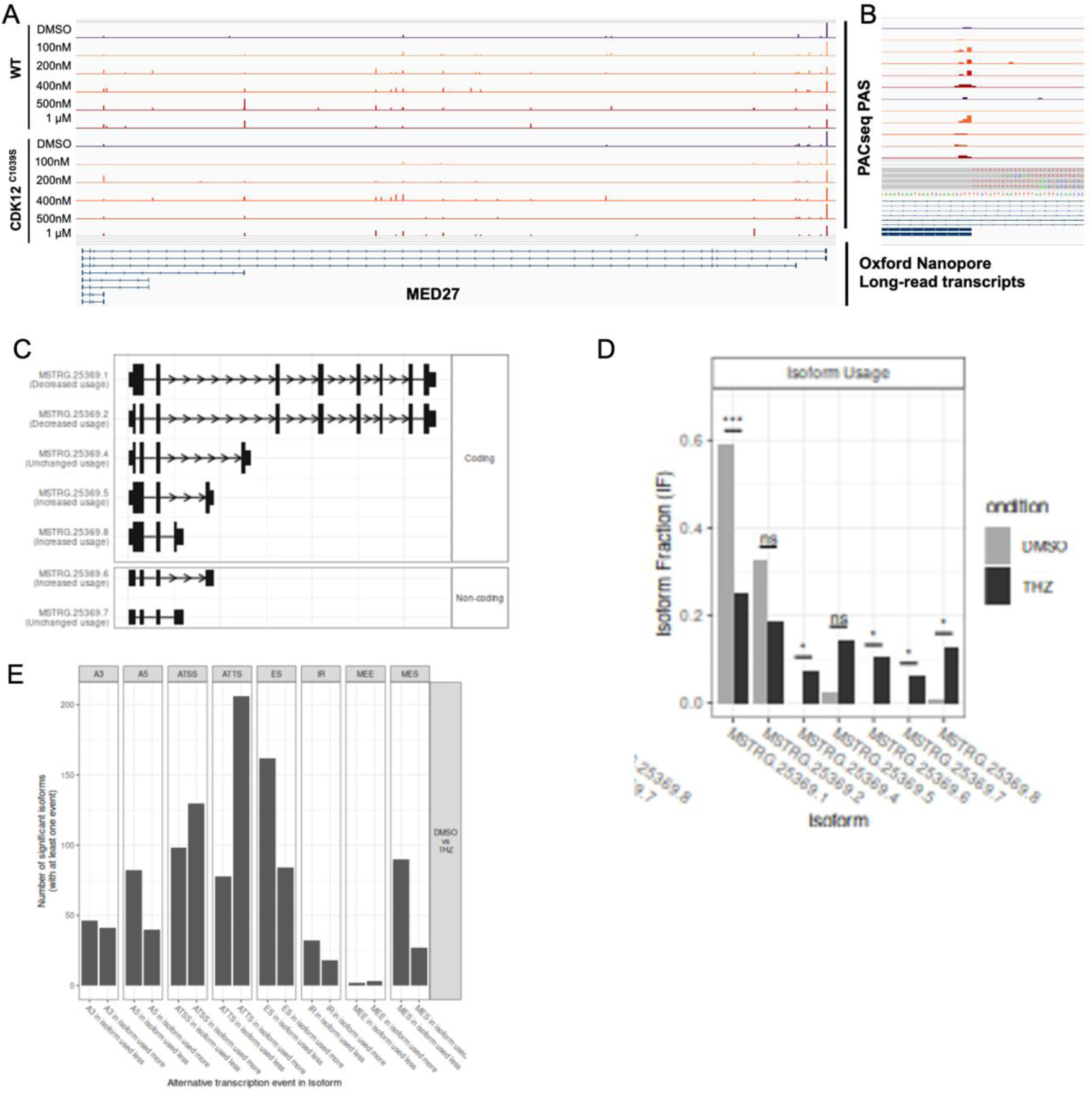
Oxford Nanopore long-read RNA sequencing identifies MED27 IPA transcript isoforms OVCAR4 WT and mutant-treated cells. A) Annotated PACseq PAS sites combined with Oxford Nanopore long-read transcripts from RNA sequencing of the transcripts MED27 in WT and CDK12 C1039S treated cells. Vertical lines on the transcript reads for each PACseq sample indicate polyA sites and the Oxford nanopore sequencing shows sequencing upstream of the polyA site and the resulting transcripts. B) An example of a single IPA site from MED27 sequencing data showing the polyA tail (polyT in cDNA/sequencing results) that is added into the prematurely terminated site within the transcript and shows IPA happens consistently at the same nucleotide base. C) IsoformSwitchAnalyzeR results of MED27 RNA sequencing data showing all peptide sequences that would arise from MED27 transcripts, including IPA transcripts. Isoforms are sorted into putative coding and non-coding sequences which would likely be degraded due to frameshifting. D) Isoform abundance of MED27 transcripts in DMSO and THZ531-treated cells. E) Genome wide identification of isoform switches, grouped by structural categories.

The implementation of long-read sequencing to determine the outcome of alternative polyadenylation events can potentially provide information that exceeds what is possible using conventional, short-read platforms such as Illumina. First, it can identify alternative start codons as well as multi-exon spanning changes in coding sequences that can profoundly affect protein function. As an example, the gene TSPAN14 contains four transmembrane domains and an extracellular domain and its altered expression due to allelic variants is associated with Alzheimer’s disease (ref). Mapping both PACseq and long-reads to the TSPAN14 gene, we were able to identify multiple IPA sites that were upregulated by THZ531 treatment (Fig. 6A). TSPAN14 is among the set of genes that contain both a significantly upregulated IPA site as well as a significantly downregulated distal UTR site based on the PACseq data. IsoformSwitchAnalyzeR identified eight isoforms, some of which were coding and some non-coding and which contained two distinct start codons and multiple termination sites resulting in different C-termini among which the isoforms terminating at the major IPA site were strongly upregulated (Fig. 6B, C). Using AlphaFold predicted structures, the consequences of the alternate start codons and termination sites were mapped onto the protein structure (Fig. 6D). Use of the internal start codon results in loss of one of the transmembrane domains and the cytoplasmic tail of the protein. Use of the IPA site results in production of a protein that is lacking a different transmembrane domain, as well as the entire extracellular domain. Combining PACseq and long-read sequencing enabled us to understand both the mechanistic underpinnings of a THZ531 sensitive isoform switch, as well as the functional consequences to protein structure in a gene with well-established importance in human disease.

**Figure 6:**
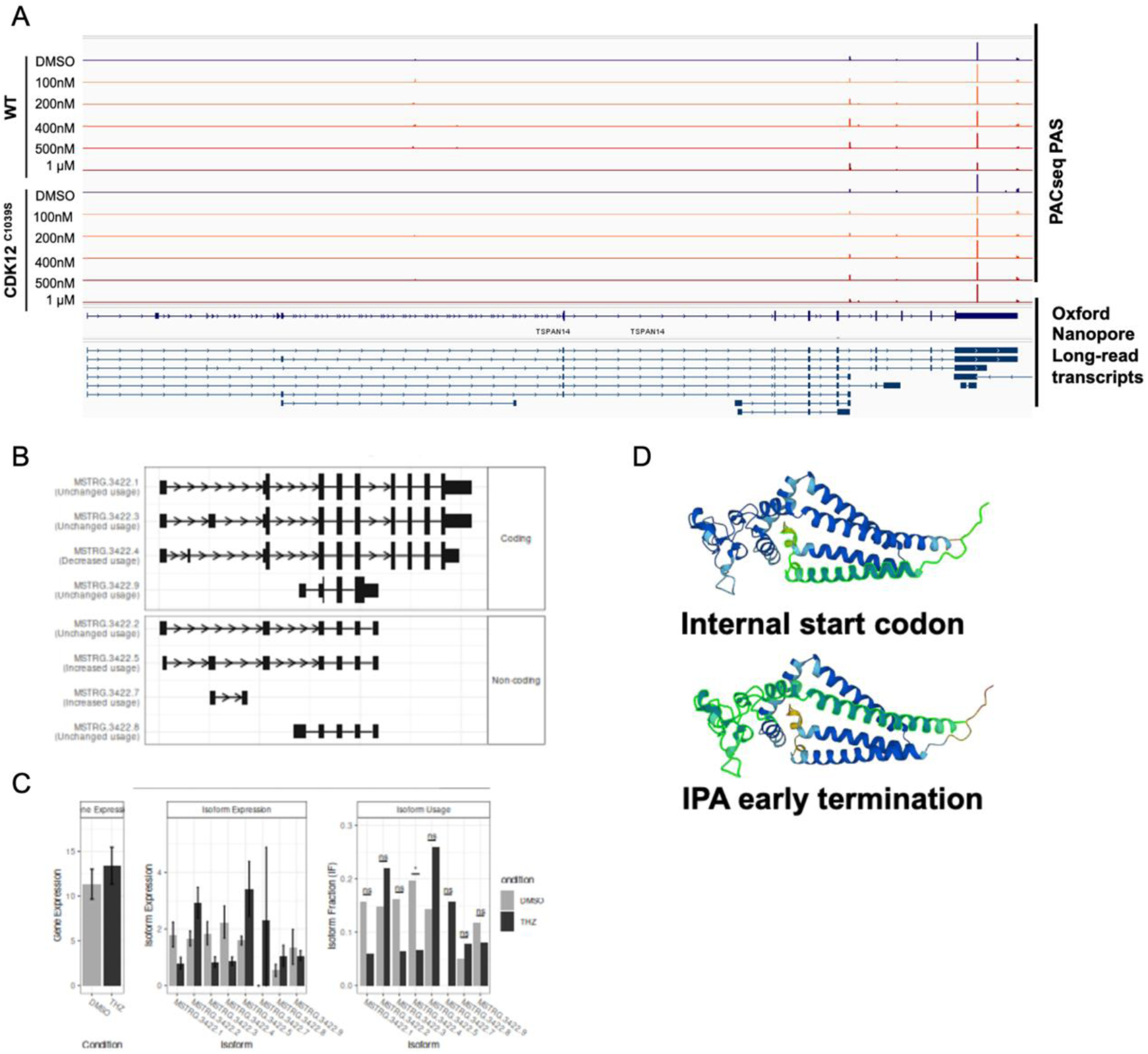
TSPAN14 is affected by IPA and translated into a truncated peptide. A) Annotated PACseq PAS sites combined with Oxford Nanopore long-read transcripts from RNA sequencing of the transcripts TSPAN14 in WT and CDK12 C1039S treated cells. Vertical lines on the transcript reads for each PACseq sample indicate polyA sites and the Oxford nanopore sequencing shows sequencing upstream of the polyA site and the resulting transcripts. B) IsoformSwitchAnalyzeR results of TSPAN14 RNA sequencing data showing all peptide sequences that would arise from MED27 transcripts, including IPA transcripts. Isoforms are sorted into putative coding and non-coding sequences which would likely be degraded. C) Gene expression, isoform expression, and isoform usage from RNA seq and proteomics data. Gene expression shows overall expression of TSPAN14 in DMSO and THZ531-treated cells. Isoform expression depicts the various transcripts that arise from different PAS usage and their relative abundance in the RNA seq data. D) Two alternate isoforms of TSPAN14 from proteomics data were mapped to AlphaFold predicted protein structure, with the part of the protein that is not present in these isoforms depicted in green.

### Proteomic analysis with expanded transcriptomes enables neopeptide identification

Previous studies have identified peptides encoded within the intronic portion upstream of an IPA site in response to CDK13 mutations^22^. It is not known whether the translation of ptRNAs produced by CDK12 inhibition also occurs. To determine whether IPA terminated transcripts produced by inhibition of either CDK12 or CDK13 are translated, we isolated cell lysates from control and THZ531-treated cells and performed whole-cell proteomics using mass spectrometry. Proper assignment of the identified peptide fragments requires a database containing the predicted sequences of the novel coding regions. To generate this database, we used our Oxford Nanopore long-read sequences compiled with IsoformSwitchAnalyzeR. An example of the problem is indicated in Fig. 7, in which the previously described IPA site isoform in TSPAN14 is shown. In transcripts terminating at an IPA, the reading frame of the immediate upstream exon continues into the intronic region until a stop codon is encountered. In TSPAN14, this results in a ten-amino acid peptide, cleaved at the exonic lysine by trypsonization, and spanning to the final arginine (Fig. 7). This peptide is detected in our proteomic data, along with several other peptides originating from IPA terminated transcripts (data not shown). Thus the early termination of transcripts at IPA sites, which occurs not only in the presence of THZ531but also in tumor cells carrying CDK12/13 mutations or mutations in other nuclear RNA surveillance genes, can produce novel peptide sequences encoded within introns. Future experiments will be required to dissect the functions of CDK12 and CDK13 in this process, since we are unable here to identify peptide in cells where only CDK12 is inhibited.

**Figure 7:**
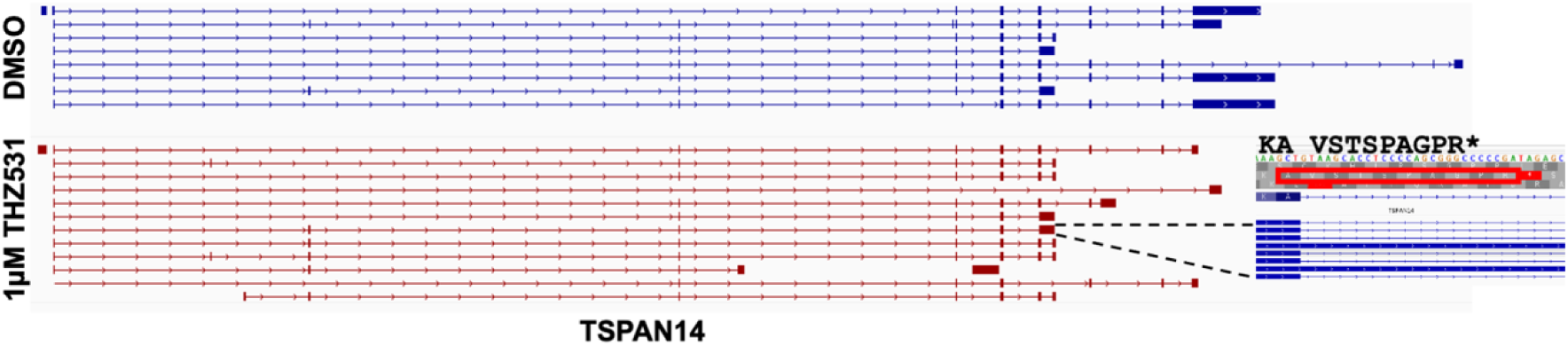
Novel intronically-encoded peptides arise in CDK12 and CDK13 LOF cells. Annotated transcript isoforms of TSPAN14 showing known exons and introns. The coding sequence is also depicted for an IPA transcript. Upstream of the IPA site the intron contains a stop codon, indicating the intronic region would be translated until reaching the stop codon.

## Discussion

CDK12 and CDK13 both display loss-of-function in various cancers that result in the same phenomenon: increased usage of intronic polyadenylation sites throughout a pre-mRNA transcript. Although CDK12 and CDK13 are both RNAPII CTD Ser2 kinases, their LOF status arise differently and in non-overlapping contexts: CDK12 undergoes bi-allelic mutations that cause the LOF phenotype in prostate, ovarian, and breast cancers whereas CDK13 has dominant negative mutations in one allele that results in LOF in melanomas, other solid tumors, and in developmental disorders^12,13,22,43^. Furthermore, in CDK13 LOF models, IPA transcripts have been shown to undergo translation into novel peptides, showcasing a potential tumor vulnerability for patients with CDK12/13 LOF cancers. If IPA transcripts are translated, they would encode an intronic region upstream of the PAS that would typically be spliced out of the mRNA transcript. These IPA-encoded peptides would therefore be unique to tumor cells with CDK12/13 LOF mutations.

Our approach was to model CDK12 LOF and CDK13 LOF separately in two engineered cell lines by taking advantage of THZ531, a dual inhibitor of CDK12 and CDK13. Unfortunately, mutating CDK13 (C1017S) to create resistance to THZ531 proved futile and we were only able to create inhibitor resistance in CDK12 (C1039S), therefore modeling CDK13 LOF. This put limitations on conclusive results but still allowed us to compare WT-treated cells, in which both CDK12 and CDK13 are inhibited, and CDK12 C1039S-treated cells, where CDK13 is inhibited but CDK12 is functional. Comparing these two data sets showed us different clusters of intronic polyadenylation sites that are differentially affected upon THZ531 treatment (Fig 2.3A and 2.3C). Furthermore, we saw that metacluster 1’s IPA sites decreased in mutant cells whereas there was little change in metacluster 2 IPA site usage in WT compared to mutant cells, suggesting metacluster 1 transcripts are predominantly CDK12-dependent and those within metacluster 2 are primarily CDK13 dependent. This further reinforces the hypothesis that CDK12 and CDK13 have unique kinase targets and/or protein associations.

More in depth sequence analysis allowed us to not only identify IPA transcripts but to visualize the transcript isoforms and their relative abundance. We see from the MED27 and TSPAN14 examples (Figs. 2.5 and 2.6) that IPA transcript isoforms are overall increased in THZ531-treated cells while there is a decrease in the full-length isoform. This aligns with both our hypothesis and previous work as we see these IPA mRNA transcripts are translated and expressed in some cases at significantly higher rates in LOF models.

The proteomic analysis offers insight into the biological impact of CDK12 and CDK13 LOF phenotypes in individuals. For example, we know one of TSPAN14’s IPA isoforms does not include one trans-membrane domain and the entirety of the protein’s extracellular domain (Fig 2.6D). Although we have not yet characterized additional isoforms in depth, we imagine TSPAN14 is one of many genes whose coding sequence is significantly affected by IPA in this manner.

One major recurrent theme in this field is that CDK12 LOF, while not directly involved in the homologous recombination pathway, phenocopies BRCAness mutations. Dubbury *et al*. showed that BRCAness genes tend to have both high levels of IPA sites and are more sensitive to CDK12 LOF than non-BRCAness genes with the same numbers of IPA sites. Combining this knowledge with our proteomic findings, there are likely a large number of BRCAness protein isoforms that have missing functional domains, as with TSPAN14.

In order to fully distinguish between CDK12 and CDK13 LOF, we will need to develop distinct LOF cell lines to model the cancer phenotypes. Once we have these systems in place, we will be able to better illustrate CDK12- and CDK13-dependent IPA events. Further proteomic analysis will lead us to identify novel intronically-encoded peptide isoforms and then to determine whether any of these neopeptides are bound to MHCI and displayed on the cell surface. Ultimately, pinpointing potential neoantigens may enable more targeted immunotherapy treatments.

## Methods

### Plasmid generation

HDR templates were cloned into the puc19 plasmid and sgRNAs were clones into the px330 plasmid. We adapted the Zhang lab target sequence cloning protocol to make px330 sgRNA plasmids. 1 µg of px330 was digested with *Bbs*I (NEB R0539S), run on an agarose gel, and purified using the QIAquick Gel Extraction Kit (Qiagen 28704). 1 µL each of sgRNA forward and reverse oligos (100 µM) were phosphorylated (T4 PNK (NEB M0201S) and annealed at 37 °C for 30 minutes and diluted 1:200. Oligos were annealed to digested and purified px330 with Quick Ligase (NEB M2200SVIAL) for 10 minutes on the bench. Assembled plasmid was then transformed with One Shot™ TOP10 Chemically Competent *E. coli* (C404010).

The puc19 plasmid was digested with *Pvu*II (NEB R0151S) and purified and phosphorylated as described with px330. 10 ng of HDR templates were phosphorylated with T4 PNK and diluted 1:50. Digested puc19 (50 ng) and the HDR template were ligated with quick ligase as described above and transformation was performed.

### Cell culture

Cell lines 22Rv1, OVCAR4, OVCAR8, and CAOV3 were obtained from American Type Culture Collection. 22Rv1, OVCAR4, and OVCAR8 cells were cultured in RPMI 1640 (Invitrogen) supplemented with 10% FBS (fetal bovine serum) and CAOV3 cells were cultured with DMEM (Invitrogen) supplemented with 10% FBS. Cell lines were maintained at 37 °C in a 5% CO_2_ cell culture incubator. To passage, cell were washed with HBSS (Invitrogen) and suspended with TrypLE Express Enzyme 1X (Invitrogen).

### Neon electroporation

The Alt-R™ CRISPR-Cas9 System from IDT was used for genome editing with electroporation. The Invitrogen Neon Electroporation System (Thermo Fisher MPK5000) was used for electroporation. Specific crRNAs were ordered from IDT along with a generic tracrRNA. 200 µM crRNA hybridized with 200 µM tracrRNA in 5.6 µL IDT duplex buffer to anneal the gRNA. Hybridization was performed by heating at 95 °C for 5 minutes then cooled on the bench for 20 minutes. Cas9 RNPs were assembled by diluting 0.3 µL of Cas9 with 0.2 µL of Neon buffer R then mixing the diluted Cas9 to 0.5 µL of the crRNA:tracrRNA gRNA. The RNP was mixed by swirling with the pipette tip to avoid introducing bubbles and then incubated for 20 minutes on the bench.

Cells were brought to a concentration of 5,000,000 cells/mL in a volume of 10 µL so that each transfection has 50,000 cells. For each transfection, the following reagents were mixed together: 1 µL of HDR oligo template; 1 µL of RNP; 3 µL of Neon buffer R; 10 µL of cells. The appropriate electroporation conditions were applied (for OVCAR4: 1170 v, 30 ms, and 2 pulses; for OVCAR8: 1600 v, 10 ms, and 3 pulses; for 22RV1: 1500 v, 20 ms, 1 pulse). Transfected cells were plated in 6-well plates. Four days post-transfection wells were expanded to 15-cm dishes.

After 2-3 weeks individual transfected clone colonies were picked from 15-cm dishes. Half of each colony was plated in a 96-well plate for expansion and the remaining half was taken for genomic sequencing by mixing with 60 µL of QuickExtract DNA Extraction Solution (Lucigen QE0905). QuickExtract-treated cells were vortexed for 15 seconds, incubated at 65 °C for 6 minutes, vortexed for 15 seconds, and incubated at 98 °C for 2 minutes. This genomic DNA was then PCR amplified, purified, and sent to Genewiz for sequencing. Sequenced and identified homozygous mutants were then expanded for further experimentation.

### Lipofectamine transfection

Lipofectamine 3000 Transfection Reagent kit (Invitrogen L3000015) was used for lipid-based transfections. Cells were plated at 100,000 cells/mL in 6-well plates 24 hours pre-transfection. 7.5 µL lipofectamine 3000 reagent was used with a total of 2500 ng DNA. 1250 ng of puc19 HDR plasmid was used with 625 ng of px330 sgRNA plasmids and 625 ng of pLKO puromycin-resistance plasmid. 24 hours post-transfection puromycin selection was performed. 72 hours post-transfection cells were expanded into 150 mm plates and grown until clone colonies were large enough to be picked and sequenced.

### THZ531 treatment

Cells were plated at 50,000 cells/mL in 15-cm dishes. After 8-12 hours either DMSO or 1mM THZ531 was added directly to the medium at a final concentration ranging between 100-1000 nM. 8 hours post THZ531 treatment, medium was removed from cells and cells were washed with cold HBSS on the bench. HBSS wash was removed and 1 mL of fresh HBSS was added to the plate. Cells were removed via scraping and volume was split three-ways; 1/3 for RNA sequencing, 1/3 for western blotting, and 1/3 for mass spectrometry.

### Protein purification, SDS-PAGE, and western blotting

Cells in HBSS were spun at 1000xg in a table-top centrifuge for 30 seconds. Supernatant was removed and cells were resuspended in 30-60 µL lysis buffer (Pierce RIPA lysis buffer (Thermo Fisher 89900); proteinase inhibitor (cOmplete™, Mini, EDTA-free Protease Inhibitor Cocktail (Sigma Aldrich #4693159001), 1 tablet/mL)); Pierce Universal nuclease (0.5 µL; Invitrogen #88701); phosphatase inhibitor (100x)). Lysis-treated cells were incubated on ice for 20 minutes and then spun at 4 °C for 10 minutes at 14,000xg. Protein-containing supernatant was saved. The Qubit Protein Assay kit (Thermo Fisher #Q33212) was used to measure protein concentration. An appropriate volume of 4X Bolt™ LDS Sample Buffer (Thermo Fisher #B0007) was added to 7.5 µg protein and incubated at 70 °C for 10 minutes. Protein was loaded in Bolt Bis-Tris Plus 4-12% (Invitrogen #NW04120BOX) and gel was run at 200 V for 22 minutes (Transfer buffer: 20X Bolt Transfer buffer #BT0006, 10% methanol). The protein was transferred from the gel to a nitrocellulose membrane for 1 hour at 70 °C at room temperature. The membrane was blocked in milk blocking buffer (50 mL TBS, 500 µL Tween20, 25 µL skim milk, to 500 mL with water) for approximately 1 hour. Blocked membranes were probed with primary antibodies overnight at 4 °C. Antibody mix was removed and membranes were washed for 5 minutes in TBS-T at room temperature 3 times. Membranes were blotted with a fluorescent secondary antibody for 1 hour and then TBS-T wash was repeated. Western blots were visualized with Typhoon FLA 9500 biomolecular imager from GE Healthcare Life Sciences.

### RNA purification and sequencing

Qiagen’s RNeasy Mini Kit (Qiagen 74106) was used to purify RNA from cells. 600 µL of buffer RLT was added to cell pellets and suspended volume was put through a QIAShredder (Qiagen 79656) to lyse cells. The RNeasy protocol was followed according to manufacturer. The Qubit™ RNA High Sensitivity kit (Thermo Fisher Q32852) was used to determine RNA concentration. Library preparation for Nanopore sequencing was performed using the PCR-cDNA barcoding kit (Oxford Nanopore #SQK-PCB111.24) with 200 ng of total RNA as the starting material. For PAC-seq, 2 µg of total RNA was sent to Azenta for library preparation and sequencing.

### Library preparation for mass spectrometry

Cells were lysed with 100 mM TEAB with 5% SDS (100 µL 1 M TEAB (Thermo #90114), 500 µL 10% SDS (Invitrogen #AM9822), 400 µL water (Fisher #W6-500)) and protein concentration was determined and 25 µg of protein was brought to a volume of 25 µL. Protein was reduced with 25 mM DTT/TEAB (100 µL 1 M TEAB, 900 µL water, 3.85 mg DTT (Thermo #R0862)) to a final concentration of 2 mM and incubated at 55 °C for 60 min at 600 rpm. Protein was alkylated with 125 mM IAA/TEAB (100 µL 1 M TEAB, 900 µL water, 23.1 mg Iodoacetamide (IAA, Thermo #AC122270250)) to a final concentration of 10 mM and incubated at room temperature for 30 min in the dark. 3.3 µL 12% phosphoric acid (142 µL 85% phosphoric acid (Fisher #A260-500), 858 µL water) was added to the protein for acidification. 6x 90% methanol/TEAB (9 mL 100% methanol (Fisher #BP1105-4), 1 mL 1 M TEAB) was added to the acidic proteins and resuspended. An S-trap column (Protifi #C02-micro) was used to bind the proteins and then the protein was washed 2 times with 90% methanol/TEAB. To digest the proteins from the column, 20 µL trypsin/TEAB (200 µL 1 M TEAB, 1800 µL water, 100 µg trypsin (Pierce #PI90058) was added and incubated for 5 minutes at room temperature. The whole column was then incubated in a 37 °C water bath for approximately 20 hours. To elute proteins, 20 µL 0.1% TFA (Thermo #LS119-500) in water was added to the S-trap and spun at 4000 xg for 1 minute. Eluate was collected. For final elution, 40 µL 50% acetonitrile in water (500 µL water, 500 µL acetonitrile (CAN with 0.1% TFA, Fisher #LS121-500)) was added and spun at 4000 xg for 2 minutes. Eluate was collected and combined with protein from the previous step.

**Table 1:**
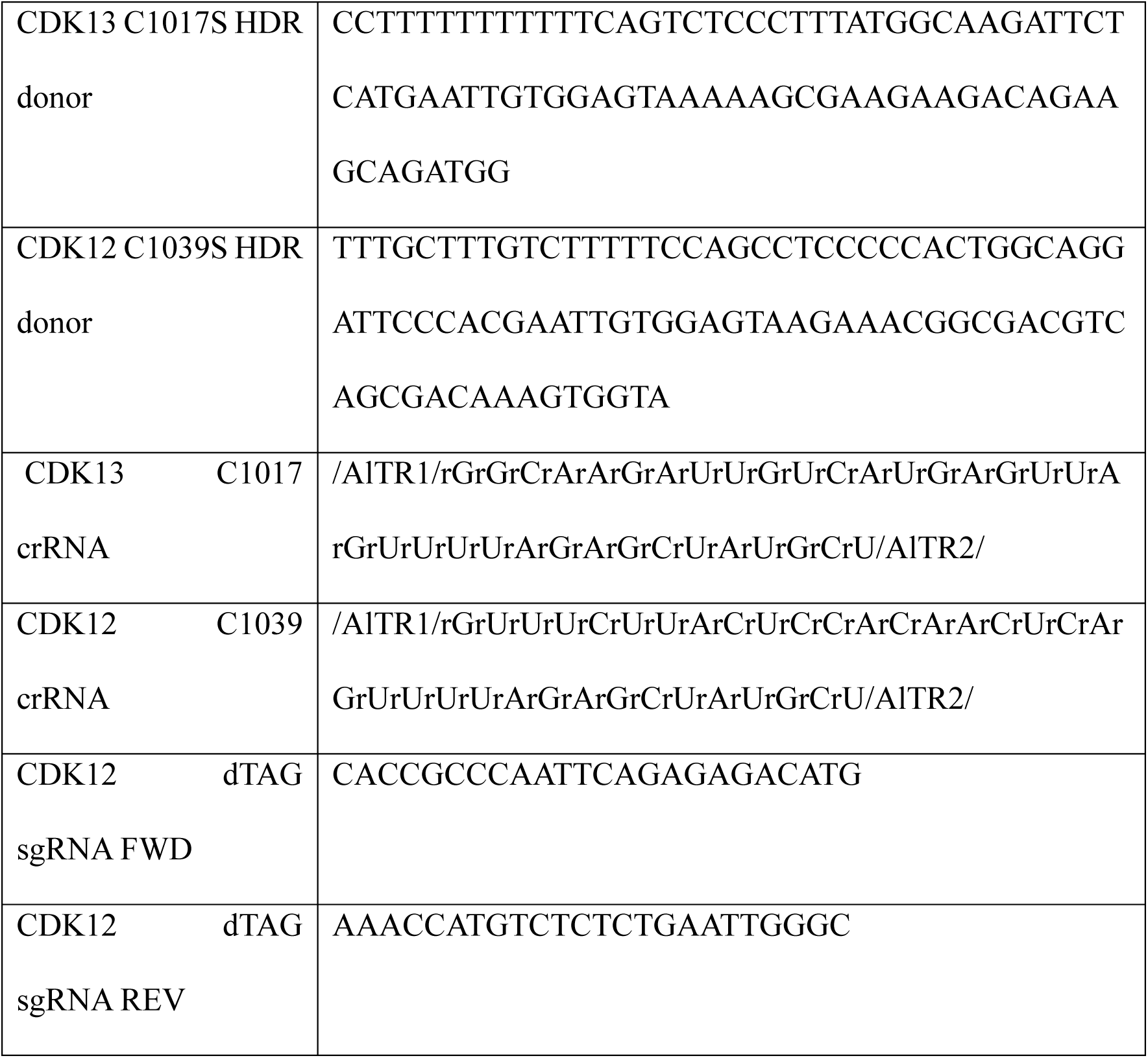
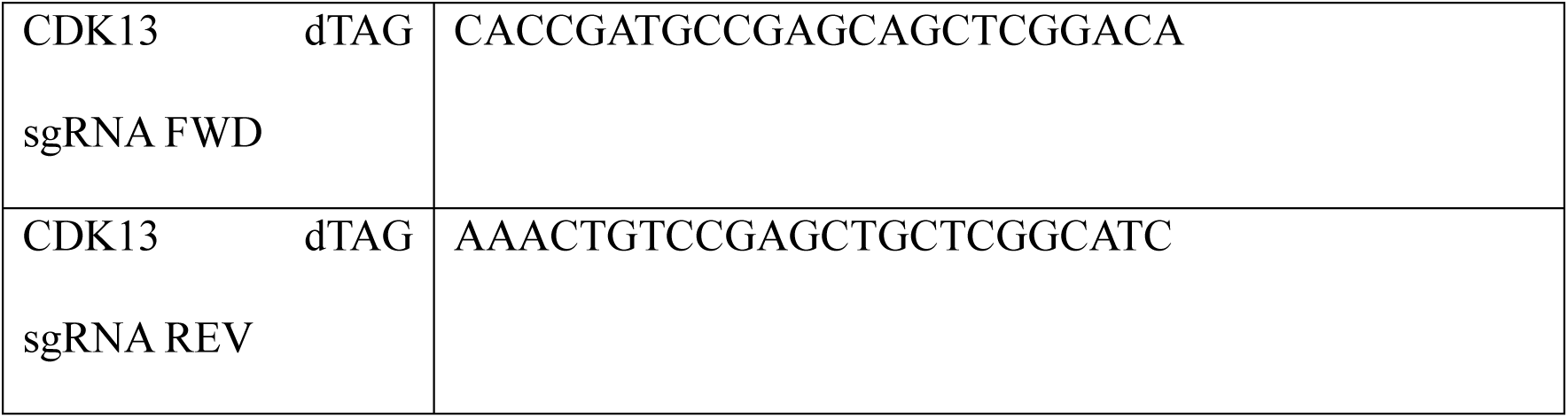
DNA and RNA oligos used for cellular transfections.

### PACseq data processing and polyadenylation site analysis

Raw Illumina sequences in the form of fastq files were first trimmed with Cutadapt (M. Martin) to remove adaptor sequences and then mapped to Hg38 using STAR version 2.5.3 (Dobin, et al. 2013). Mapped reads were then processed using a custom computational pipeline. Briefly, mapped reads were first filtered to require a minimum of 6 nucleotides at the 3’ end of the read (based on transcriptional direction) that are soft-clipped, that is, that don’t align to the contiguous genomic sequence. These 6 nucleotides were further required to be only adenosine to ensure a poly(A) tail is present. Using the CIGAR field as a unique molecular identifier, duplicate reads were also discarded at this step. Reads were then grouped by biological treatment, and filtered to ensure the presence of a canonical polyA signal sequence (AATAAA, ATTAAA, or one of 6 other minor variants) within the -10 to -40 region upstream of the cleavage site. Reads were assigned to annotated genes, and sites overlapping coding exons were eliminated.

Cleavage sites falling within 40 nucleotides of each other were clustered together using bedtools (Quinlan et al, 2012) and custom python scripts. After all clusters from individual treatments were clustered, the entire data set was combined for reclustering, using a minimum threshold abundance level corresponding to 0.1% of the cleavage sites within a given gene to remove low-end noise. This resulted in the final set of 256,498 PAS clusters. To perform quantitative analysis, single-nucleotide bams were generated that included only the 3’ most genomically mapped nucleotide (corresponding to the last nucleotide prior to the poly(A) tail). We then determined the number of read counts falling within each PAS cluster. These raw counts were then used in DESeq2 to identify, genome-wide, the significantly affected sites (p adj < 0.05) when comparing the DMSO vs. 100nM treatments, or a combination of the the 500nM and 1uM treatments.

To generate heat maps, the fraction of transcripts using each PAS in every replicate was determined. The fractions were averaged across replicates of the same treatment and plotted on a color scale ranging from 0 to 1, with 0 being no usage and 1 being 100% of the transcripts in the gene. These were then hierarchically clustered using the differentials at each dose to generate clusters and metaclusters. Dose curves were then fitted to each group based on all of the fraction numbers within the dosages. Venn diagrams were produced using Venny2.0.

### Oxford Nanopore Long-read Sequencing and analysis

Libraries were created using the Oxford Nanopore cDNA barcoding kit according to manufacturer’s instructions. Sequencing was performed on a Promethion P2 Solo device. Reads were mapped using Epi2Me, minimap2, and Stringtie. Long read isoform switching data was analyzed in R using the package IsoformSwitchAnalyzeR.

